# The Recombinational Landscape in *Daphnia pulex*

**DOI:** 10.1101/2020.03.03.974485

**Authors:** Michael Lynch, Zhiqiang Ye, Takahiro Maruki

**Affiliations:** Biodesign Center for Mechanisms of Evolution, Arizona State University, Tempe, AZ 85287

## Abstract

Through the analysis of linkage disequilibrium from genome-wide sequencing data for multiple individuals from eight populations, the general features of the recombinational landscape are revealed in the microcrustacean *Daphnia pulex.* The data suggest an exceptionally uniform pattern of recombination across the genome, while also confirming general patterns that are inconsistent with existing population-genetic models for the relationship between linkage disequilibrium and physical distances between genomic sites. Patterns of linkage disequilibrium are highly consistent among populations, and average rates of recombination are quite similar for all chromosomes. There is no evidence of recombination hotspots, and although there does appear to be suppressed recombination in the vicinity of gene bodies, this effect is quite small. Although this species reproduces asexually in ∼ 80% of generations, the mean per-generation recombination rate per nucleotide site is ∼ 37× the per-nucleotide mutation rate. Contrary to expectations for models in which crossing-over is the primary mechanism of recombination, and consistent with data for other species, the gradient of linkage disequilibrium with increasing physical distance between sites is far too high at short distances and far too low at long distances, suggesting an important role for factors such as the nonindependent appearance of pairs of mutations on haplotypes and long-range gene-conversion-like processes. Combined with other observations on patterns of nucleotide variation, these results provide a strong case for the utility of *D. pulex* as a model system for the study of mechanisms of evolution in natural populations.

---

Recombination plays numerous roles in evolutionary processes, providing a path to the joint appearance of independent mutations in the same haplotype while also freeing beneficial mutations from background deleterious mutations at linked sites (Charlesworth and Charlesworth 2010). Despite these presumed advantages of recombination, one of the most conserved genetic features across the entire eukaryotic domain is the occurrence of just one to two crossovers per chromosome arm per meiotic event, which results in an inverse relationship between the crossover rate per nucleotide site and genome size (Lynch 2007; Lynch et al. 2011). In principle, recombination rates between closely spaced sites can be much higher than expectations based on crossovers if a substantial fraction of recombination events involves gene-conversion events with nonreciprocal exchange, as seems to be the case (Andolfatto and Nordborg 1998; Lynch et al. 2014). Moreover, in some mammals recombination events are highly concentrated into relatively small fractions of the genome, such that some genes experience much higher rates of both gene conversion and crossing over than others (Auton et al. 2012).

The latter types of effects (gene conversions and recombinational hotspots) are not easily revealed by conventional meiotic genetic maps, which generally capture only crossover events and typically not very many of them. However, patterns of linkage disequilibrium in samples from natural populations have the potential to illuminate fine-scale recombinational features with cumulative effects ranging over time scales roughly equivalent to the effective population size. As population-genomic data become increasingly common, we can therefore expect such analyses to greatly refine our understanding of the recombinational landscape and its degree of heterogeneity across chromosomal regions and among species.

Here, in one of the largest projects of this sort ever performed, we take advantage of substantial population-genomic data sets derived from several populations of the aquatic microcrustacean *Daphnia pulex* to reveal the basic recombinational features in this model species. The data indicate a relatively uniform recombinational landscape across the genome of this species, both among populations and among chromosomes, with no evidence of recombination hotspots. As is becoming increasingly clear in other species, the vast majority of recombination events occur without crossing over, and a large fraction of these events seem to involve relatively long tracts of gene-conversion-like processes.

## Materials and Methods

### Sample collection and DNA sequencing

Population samples were collected in the spring of 2013 or 2014 from eight temporary-pond populations distributed in the midwest and eastern portions of North America (Maruki et al., in prep.) As in our prior work (Lynch et al. 2017), to maximize the likelihood that each isolate would represent a hatchling from a unique resting egg, adults were collected from the water column before reproduction had occurred. Such hatchlings would have resulted from a mating in the immediately preceding year, or possibly earlier, as resting eggs can survive multiple years under certain conditions (although long-term survival seems unlikely in highly oxidized surface soil). Consistent with most recruitment having derived from the most recent bout of sexual reproduction, all population samples were in near Hardy-Weinberg equilibrium proportions (Maruki et al., in prep.) Clones were expanded in the laboratory, DNA extracted, and sequenced and screened as previously outlined (Lynch et al. 2017).

### Read mapping and data filtering

All sequence reads were mapped to the reference genome (PA42; Ye et al. 2017), derived from a midwest US clone of *D. pulex*, again as outlined in Lynch et al. (2017). As more fully detailed in Maruki et al. (in prep.), the resultant clone-specific data within each population were then screened to mask out any sites discordant with a goodness-of-fit test, to remove any potentially contaminated isolates and/or members of pairs of close relatives, and to remove clones with < 3× average coverage per site. The resultant sample sizes for populations, which range from 71 to 93, are provided in Maruki et al. (in prep.)

To minimize potential issues with mismappings of reads to paralogous or repetitive regions and to avoid sites subject to high error rates, based on the overall distribution of coverage across the genome, we further restricted all analyses to sites having between ∼ 50 and 150% of the modal population-level coverage across the genome within each population as well as to sites with error-rate estimates < 0.01 (see Lynch et al. 2017). The resultant numbers of sites analyzed per population are noted in Maruki et al. (in prep.). For fine-scale linkage-disequilibrium analyses, we also ignored all pairs of sites separated by gaps containing > 10 nucleotide sites, as well as any sites within 1-kb flanking regions of such regions.

### Estimation of broad-scale patterns of linkage disequilibrium

Site-specific allele and genotype frequencies, and individual genotype calls were determined using the maximum-likelihood (ML) procedures of Maruki and Lynch (2015, 2017). Population-level linkage disequilibrium was estimated using the ML method of Maruki and Lynch (2014), splitting scaffolds at any locations containing gaps ≥ 10 nucleotides. Individual-level disequilibrium (correlation of zygosity) was estimated with the methods of Lynch (2008) as implemented in Haubold et al. (2010). These methods are easy to implement, requiring as input the simple quartets of reads (numbers of inferred A, C, G, and T nucleotides) at each site in each individual, and factor out all sources of sequencing errors (not simply quality scores). For the mean sequence coverages and numbers of individuals deployed in this study, these methods yield estimates that are essentially unbiased with sampling variance close to the minimum theoretical possibility, and hence perform as well as (and in many cases better than) other existing methods (Maruki and Lynch 2015, 2017).

To obtain individual-level LD estimates, results for each population were averaged for each of the ten individuals with the highest sequence coverage. The correlation of zygosity was obtained for sites separated by distances 1 to 250,000 bp for each clone. For the first 1000 bp, estimates were obtained for increments of single nucleotides, whereas from 1000 bp to 20,000 bp, the data were binned into windows 10-bp in width, and for larger distances into 100-bp bins.

### Fine-scale patterns of linkage disequilibrium

To determine the average chromosomal patterns of the population recombination rate per generation per base pair (*ρ* = 4*N*_*e*_*c*, where *N*_*e*_ and *c* are the effective population size and recombination rate per generation per base pair) in each population, we attempted to estimate spatial patterns of *ρ* across the genome by two different methods. First, we applied the LDhat program of McVean and Auton (2007) to each individual population, averaging the site-specific measures across all populations to minimize background sampling noise and provide the species-wide pattern.

Second, we estimated the correlation *r*^2^ between pairs of SNPs using the maximum-likelihood (ML) method of Maruki and Lynch (2014) in sliding windows containing 101 SNPs with 21 SNPs overlapping between adjacent windows (restricting analyses to sites with minor-allele frequencies deemed to be significant at the 0.05 probability level). As discussed in the text, the upper bounds of *r*^2^ estimates are mathematically constrained by the SNP frequencies at the constituent sites, and to avoid the resultant interpretative problems, in some applications each estimate of *r*^2^ was standardized by dividing by its maximum value (conditional on SNP frequencies), yielding a parameter estimate 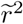 that covers the full (0,1) range. As described in the text, each of these standardized values was then converted to an expected value of *ρ*, denoted as *ρ*^*^, with the full set of such estimates within each window then being combined into an arithmetic mean for each population. Again, the population-specific values were averaged to obtain the overall species-wide pattern. Although these two approaches yield correlated results, the absolute magnitudes of the estimates of *ρ* are quite different, likely for reasons outlined in the text, with the values for LDhat being suspiciously low.

## Results

### Broad-scale Patterns

The average scaling of LD with physical distance between sites was evaluated in two ways. First, population-level LD was quantified with the squared correlation (*r*^2^) involving the joint distribution of pairs of SNPs across individuals within populations. This conventional measure of LD is not ideal, in that the upper bounds of possible values depend on the underlying allele frequencies, with maximum values of 1.0 only possible when allele frequencies at both sites are identical (VanLiere and Rosenberg 2008). The latter condition is approached as analyses are confined to specific bins of allele-frequency classes, but is still not entirely satisfied even if the analysis is restricted to very high-frequency alleles.

As the pattern of LD is quite similar among populations (below), the results are simply summarized by averaging over all eight populations for five sets of analyses with different bins for the range of minor-allele frequencies (MAFs) (Figure 1). As expected, the heights of the curves decline with decreasing MAFs. However, all five analyses exhibit consistent behavior. For sites separated by *d* ≤ 30 bp, there is an exponential decline in *r*^2^ with increasing physical distance between sites (revealed as a linear regression on a log-log plot), but for *d* > 30 bp, there is a slow quadratically increasing rate of decline in *r*^2^. Together, these two fits almost perfectly describe the scaling behavior of average *r*^2^ values within all classes of MAFs (Table 1). With the large numbers of genotypes analyzed in this study, even at a distance of 250 kb (the maximum to which we extended the analyses), LD is still discernible, although the values are very low.

**Table 1.**
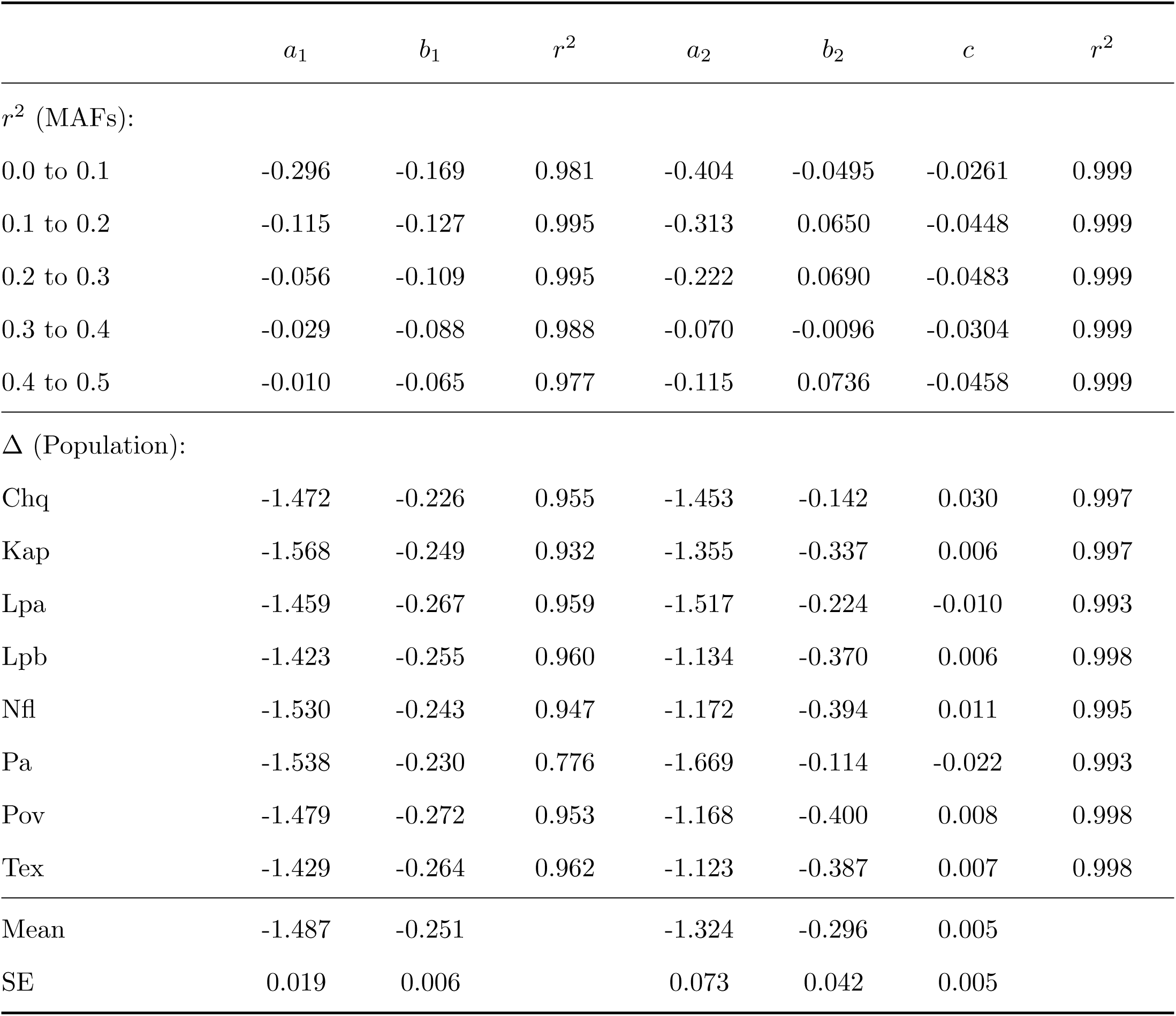
Scaling parameters for the decline of linkage-disequilibrium with physical distance *d* (in bp) between sites. Least-squares fit parameters are given for two regressions involving population-level *r*^2^: *y* = *a*_1_ + *b*_1_*x* for *d* ≤ 30; and *y* = *a*_2_ + *b*_2_*x* + *cx*^2^ for *d* > 30, where *y* = log_10_(*r*^2^) and *x* = log_10_(*d*), with results given for analyses restricted to sites with minorallele frequencies (MAFs) within five ranges.

**Figure 1.**
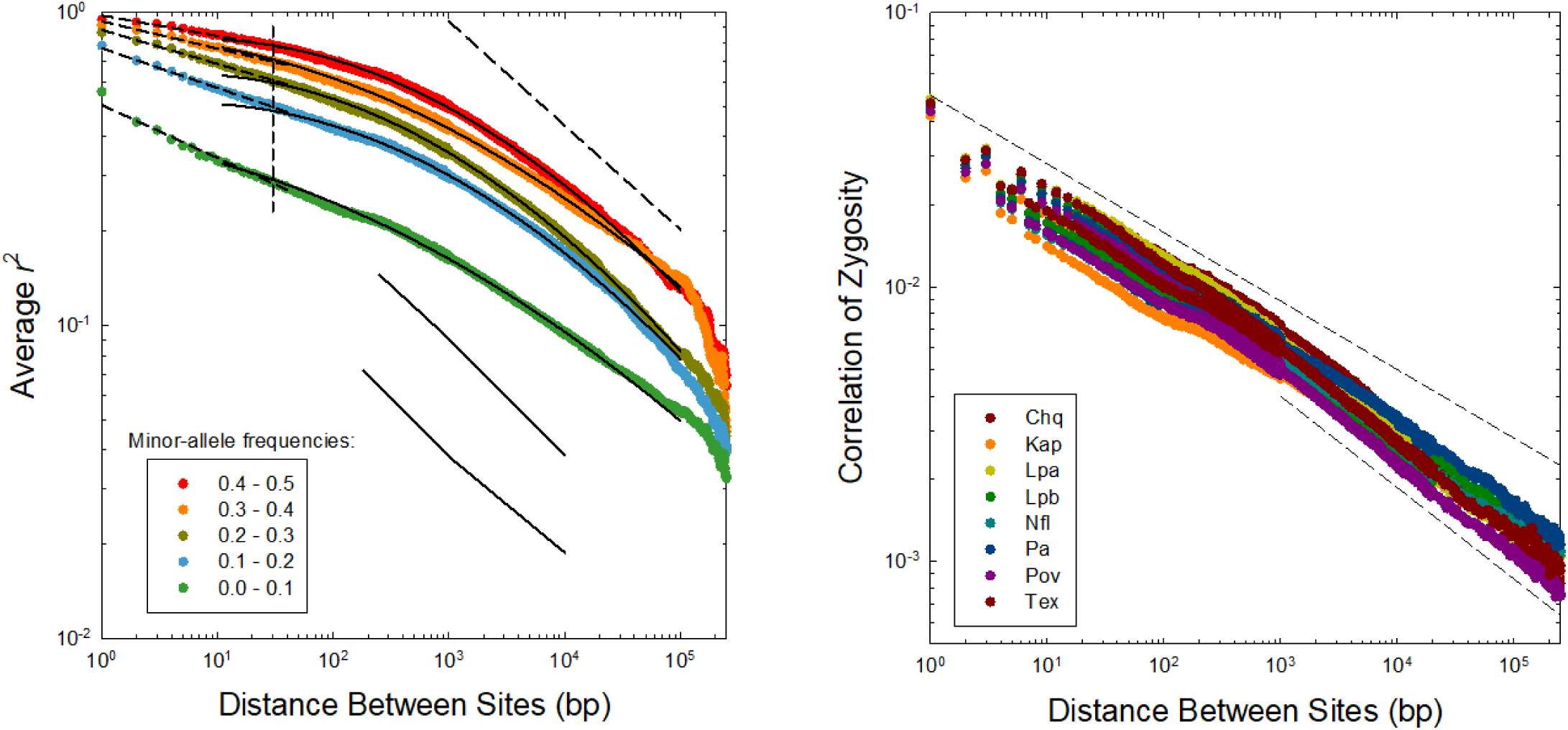
**Left)** Response of average population-level r^2^ to increasing distance between sites. Fitted least-squares regression parameters above and below this threshold are given in Table 1. The vertical dashed line at *d* = 30 bp denotes the approximate break point in the scaling behavior of LD with physical distance between sites, and the diagonal dashed line denotes −1/3 power-law scaling. The lower, thin lines give the average estimates of r^2^ for *D. melanogaster* taken from Figure 1 in Garud and Petrov (2016); upper and lower lines denote data for samples from Raleigh and Zambia. **Right)** Response of individual-level Δ to increasing distance between sites for the eight study populations. Upper and lower dashed lines denote power-law scaling behavior with slopes of −1/4 and −1/3.

Second, the correlation of zygosity, Δ, provides an individual-based measure of LD, defined by the spatial distribution of pairs of heterozygous sites. The decline of Δ with physical distance among sites is qualitatively similar among all populations, with the full range of values at any particular distance varying by a factor of no more than two between populations (Figures 1 and 2). Again, it can be seen that the distance-dependence of this measure of LD is best described by two phases – on a logarithmic scale, an approximately linear decline of Δ with distance between sites (*d*) < 1 kb, and weakly quadratic behavior at larger distances (Table 1).

**Figure 2.**
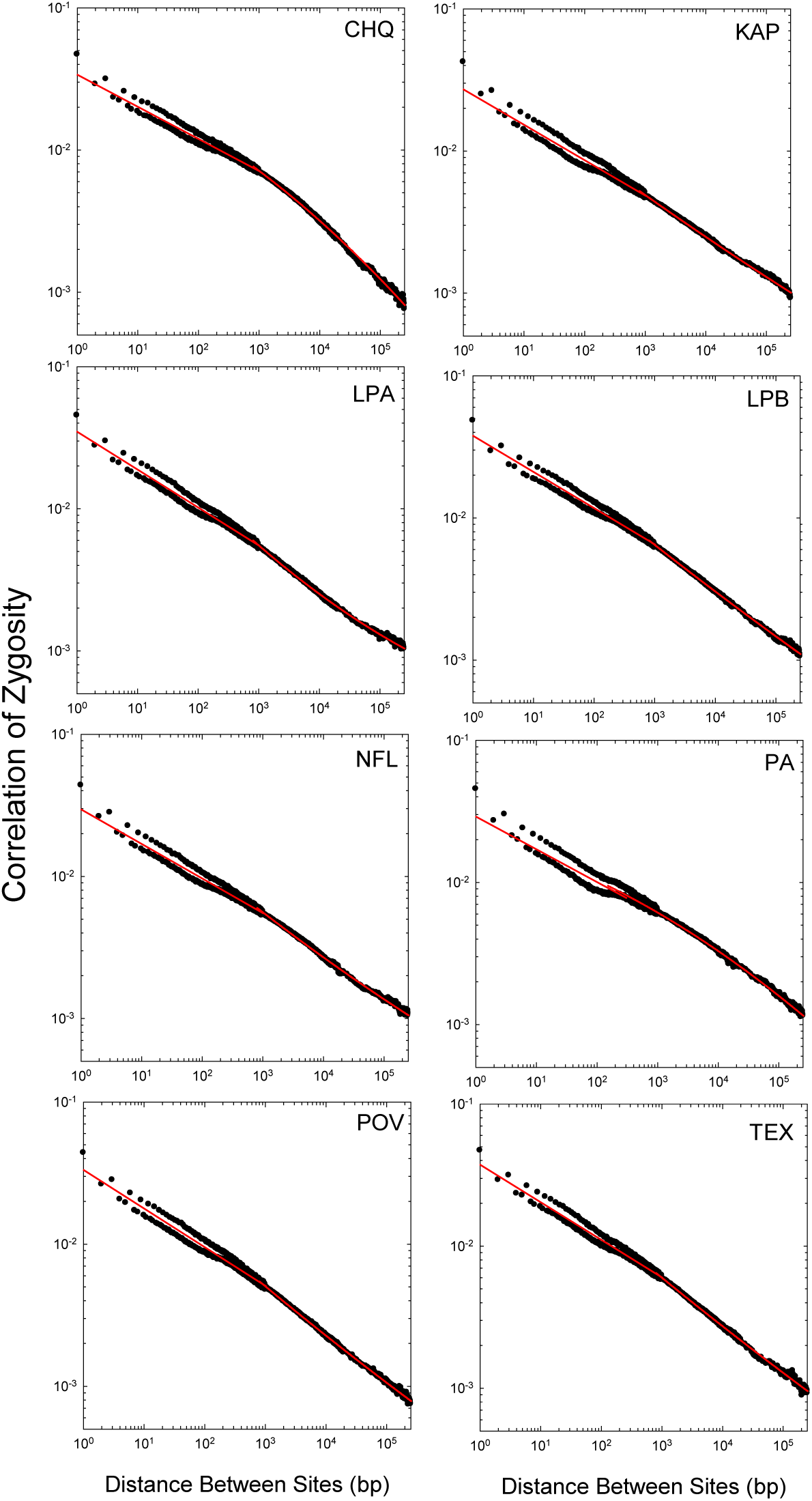
Population-specific pattern of distance-dependence of the Δ measure of individual-based LD. Red lines denote the regressions given in Table 1.

As noted in Lynch et al. (2014), Δ and *r*^2^ are expected to exhibit similar scaling with *d* under the assumption of a population in drift-mutation-recombination equilibrium, as both are proportional to the squared disequilibrium coefficient. Although average *r*^2^ scales more weakly with *d* than does Δ for *d* < 1000 bp, this is a natural artifact of the former being mathematically constrained by allele frequencies, whereas Δ is not. The vast majority of polymorphic sites have minor-allele frequencies (MAFs) ≪ 0.1 (Maruki et al., in prep.), and as can be seen in Figure 1, as the utilized allele frequencies approach zero, the scaling of *r*^2^ with distance becomes more similar to that for Δ, as expected. As a first-order approximation, for both average *r*^2^ and Δ, the negative scaling of LD with physical distance approaches a power-law relationship with exponent ≃ −1*/*4, i.e., an ∼ 1.8-fold decline in LD with a 10-fold increase in *d*, far from the expectation if the recombination rate between sites were to increase linearly with *d*.

These scaling patterns of the decline of LD with physical distance, most notably the relatively slow decline even at physical distances exceeding 1 kb, are observed in many other animals, including *Drosophila* (Lynch et al. 2014, 2017). As can be seen in Figure 1, the negative scaling of *r*^2^ with physical distance in both the North American and Zambian populations of *D. melanogaster* is similar in the range of 10^2^ and 10^4^ bp to that for *D. pulex*, although the absolute values of average *r*^2^ differ (owing in part to different ranges of allele frequencies applied).

Under drift-mutation-recombination equilibrium and the assumption that all recombination is a consequence of crossing over, the expected value of *r*^2^ is inversely proportional to *ϕ* + 4*N*_*e*_*cd*, where *ϕ* is a constant (between 1 and 4) that depends on the standing level of variation and allele frequencies, *N*_*e*_ is the effective population size, and *c* is the recombination rate per nucleotide site (Ohta and Kimura 1971; Hill 1975). Generously allowing *ϕ* = 4, based on arguments made below, this implies that if crossing over were the predominant mode of recombination, *r*^2^ would scale very nearly as 1*/*(0.05*d*) provided *d* > 200 bp. However, as just noted, even at *d* > 100 kb, the negative scaling of both Δ and *r*^2^ with *d* is far weaker than this expectation.

These observed distant-dependent patterns of LD cannot be easily resolved with conventional models for the recombination process. Under virtually all existing population-genetic models, LD is expected to be nearly unresponsive to distances at the shortest scales, as the likelihood of any recombination between sites is then very low relative to the power of drift. Yet as can be seen in Figures 1 and 2, the negative scaling of LD with distance on a scale of 1 to 100 bp is very similar to that on a scale of 100 to 1000 bp. As reviewed previously (Lynch et al. 2014), the excess LD at short distances is likely associated with the simultaneous appearance of multiple mutations on short spatial scales, which serves as a recurrent input of new LD.

High rates of gene conversion unaccompanied by crossing over can yield shallower patterns of decline of LD than expected under a model in which all recombination involves crossing over (Andolfatto and Nordborg 1998), but only at certain scales of distance. Assuming an exponential distribution of conversion tract-lengths with mean *L*, the recombination rate between sites is expected to scale as *cd* at distances ≪ *L*, with *cxd* at distances ≫ *L*, where *x* is the fraction of recombination events accompanied by crossing over, and with a flatter scaling in the transition between these domains (Figure 3). However, the scaling pattern observed in *Daphnia* cannot be fully accommodated by this model. Considering the full range of values of *x* and *L* inferred from prior work with animals (summarized in Lynch et al. 2014), low values of *x* and/or long average conversion-tract lengths (on the order of 10 kb) cause the first inverse-scaling relationship to emerge too early to be consistent with the data, whereas large *x* and/or short lengths (1000 bp) cause the second inverse relationship to appear too early.

**Figure 3.**
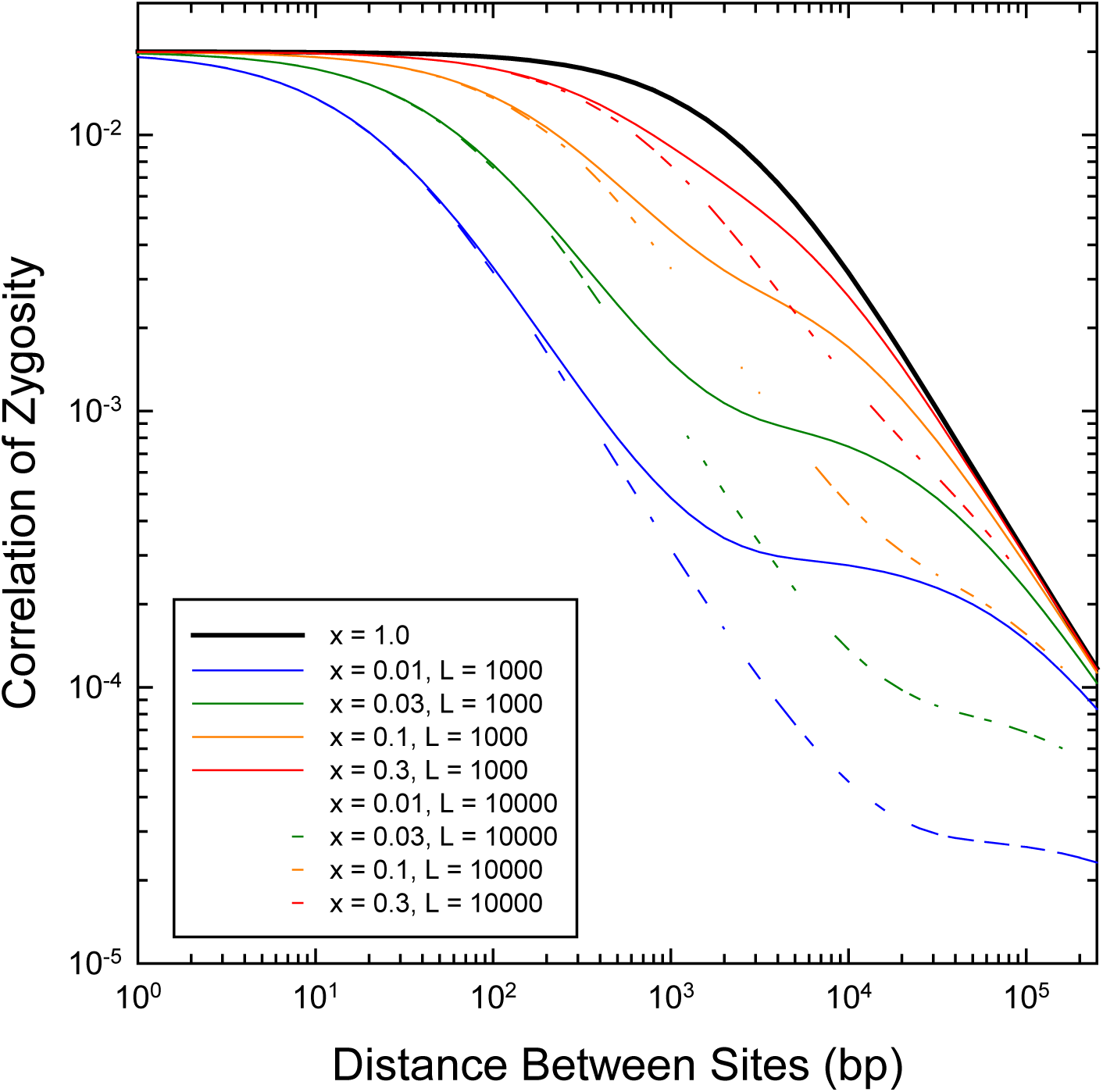
Expected pattern of decline of the correlation of zygosity with increasing physical distance between sites as a function of *x*, the fraction of recombination event resulting in a crossover and *L*, the mean length of a gene-conversion tract, under the assumption of an exponential length distribution of the latter. Derived using Equations (8) and (13a) in Lynch et al. (2014).

A likely explanation for these discrepancies is that the lengths of gene conversions (or other homogenizing conversion-like mechanisms) are not adequately described by an exponential distribution. By observing the loss of heterozygosity in asexually propagated lines, Omilian et al. (2006) and Keith et al. (2016) found that nonmeiotic homogenizing events occur in *D. pulex* at rates on the order of 10^−6^ to 10^−4^ per nucleotide site per generation, with many spans with lengths in the range of 100 to 1000 bp (as expected for gene-conversion events) but others extending up to 1000 kb (and probably involving other types of events, such as break-induced replication). In principle, a substantial frequency of long conversion-like events could retain LD over unusually long distances as pairs of markers within such spans will have their phases unaltered.

### Fine-scale Patterns

To obtain an approximate view of the landscape of recombination intensity along chromosomal regions, we employed LDhat (McVean and Auton 2007) to estimate the spatial pattern of *ρ* = 4*N*_*e*_*c* across assembled scaffolds. Given the preceding results, we cannot expect this approach to yield unbiased estimates of region-specific recombination rates, as the general procedure assumes a linear relationship between the recombination rate and the distance between sites. However, there is expected to be a monotonic relationship, on average, between the estimate 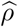 and the population parameter 4*N*_*e*_*c*. As can be seen in Figure 4 for the largest *D. pulex* scaffold, there is an approximate order-of-magnitude range in *ρ* estimates among sites, with considerable correlation in estimates from different populations.

**Figure 4.**
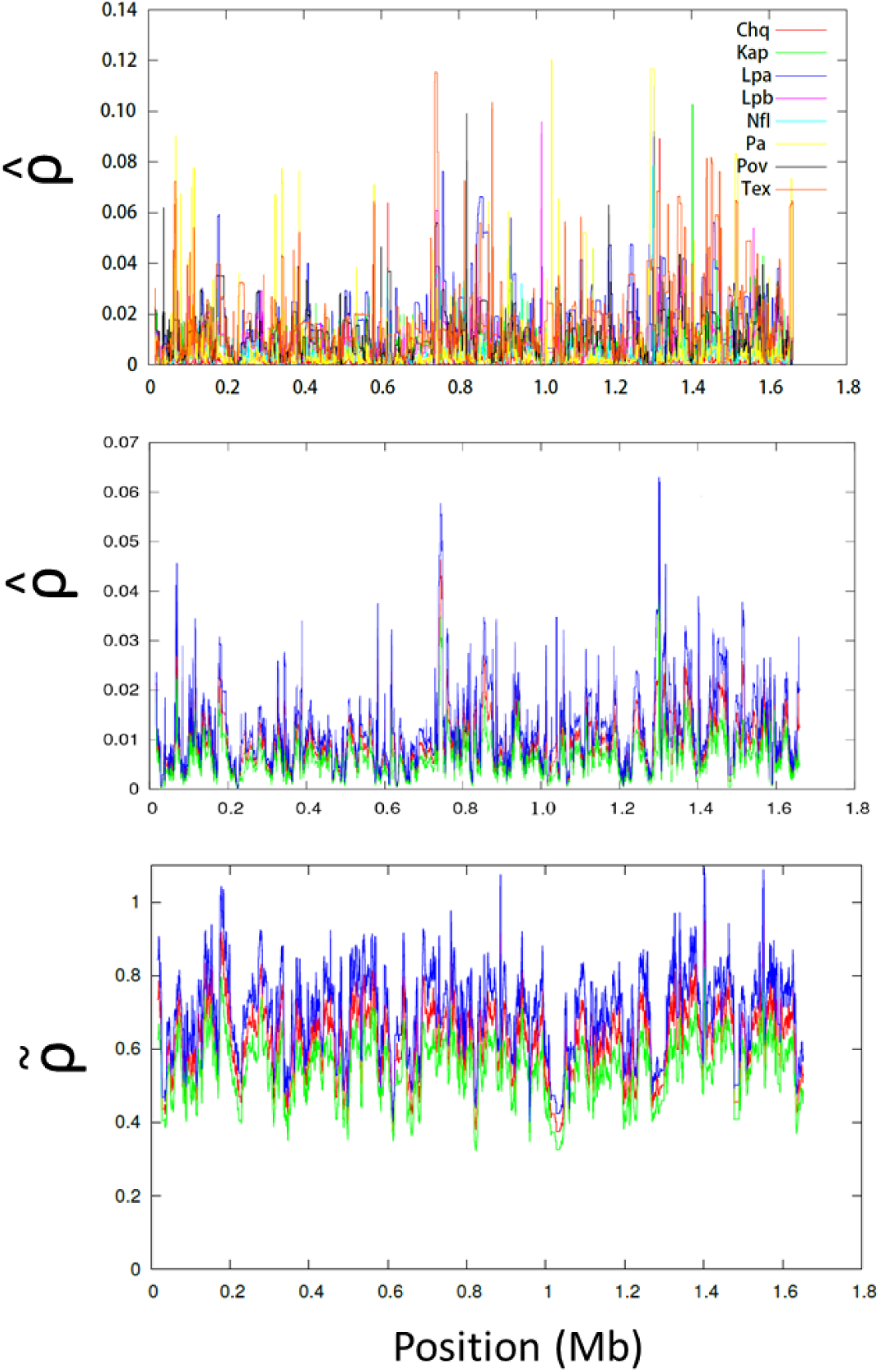
Estimated pattern of recombination-rate variability over the length of the largest *D. pulex* scaffold. The top panel overlays the 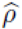 profiles for each of the eight study populations, whereas the middle panel gives the mean (red) and ± 1 one standard error (blue and green) over all populations. The lower panel gives the parallel data for 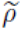.

With estimates of 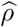 available for eight populations, it was possible to estimate the meta-population-wide mean and SEs of the population recombination rate over all genomic sites. The mean estimate of 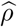 per nucleotide site is 0.0124 (SE < 10^−6^ based on 2-kb nonoverlapping windows), and the amplitude of spatial fluctuations is substantially suppressed as the noise in population-specific estimates is averaged out. 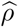 is expected to be an underestimate of 4*N*_*e*_*c* owing to the much slower rate of decline of LD than expected under the inverse-scaling relationship. However, we can more directly estimate the true value of 4*N*_*e*_*xc* (the amount of recombination associated with crossovers) by noting from the genetic map in *D. pulex* that *xc* ≃ 1.6 × 10^−8^ per nucleotide site (Xu et al. 2015), with *x* being the fraction of recombination events resulting in a crossover. With the added estimate of *N*_*e*_ ≃ 8 × 10^5^ (Lynch et al. 2017), this implies 4*N*_*e*_*xc* ≃ 0.05, and if *x* ≃ 0.1, as is the case in *Drosophila*, humans, and land plants (reviewed in Lynch et al. 2014), then 4*N*_*e*_*c* ≃ 0.5. Thus, presumably by not accounting for the nonlinear relationship between the recombination rate and physical distance, LDhat underestimates the recombination rate due to crossovers alone by 3.5× and the total rate of recombination per site (including gene conversions unaccompanied by crossing over) by 35×.

In principle, although 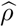 is a scaled down estimate of true *ρ*, variation in the former should still provide insight into differences in recombinational activity across chromosomal regions. The distributions of the site-specific rate estimates (as well as those for 2-kb windows) are nearly exponential (Figure 5). Although some recombination hotspots might be present, they do not account for much of total recombination. Rather, 20% of the total recombination is accounted for by ∼ 5% of the sites (or windows), 50% by ∼ 20% of the sites, and 90% by ∼ 65% of the sites. This is a strong contrast with the situation in humans, where ∼ 85% of total recombination is accounted for by ∼ 10% of the sites (Auton et al. 2012). Indeed, there is less variance in the *D. pulex* recombinational landscape than any other species for which such data are available (Figure 5).

**Figure 5.**
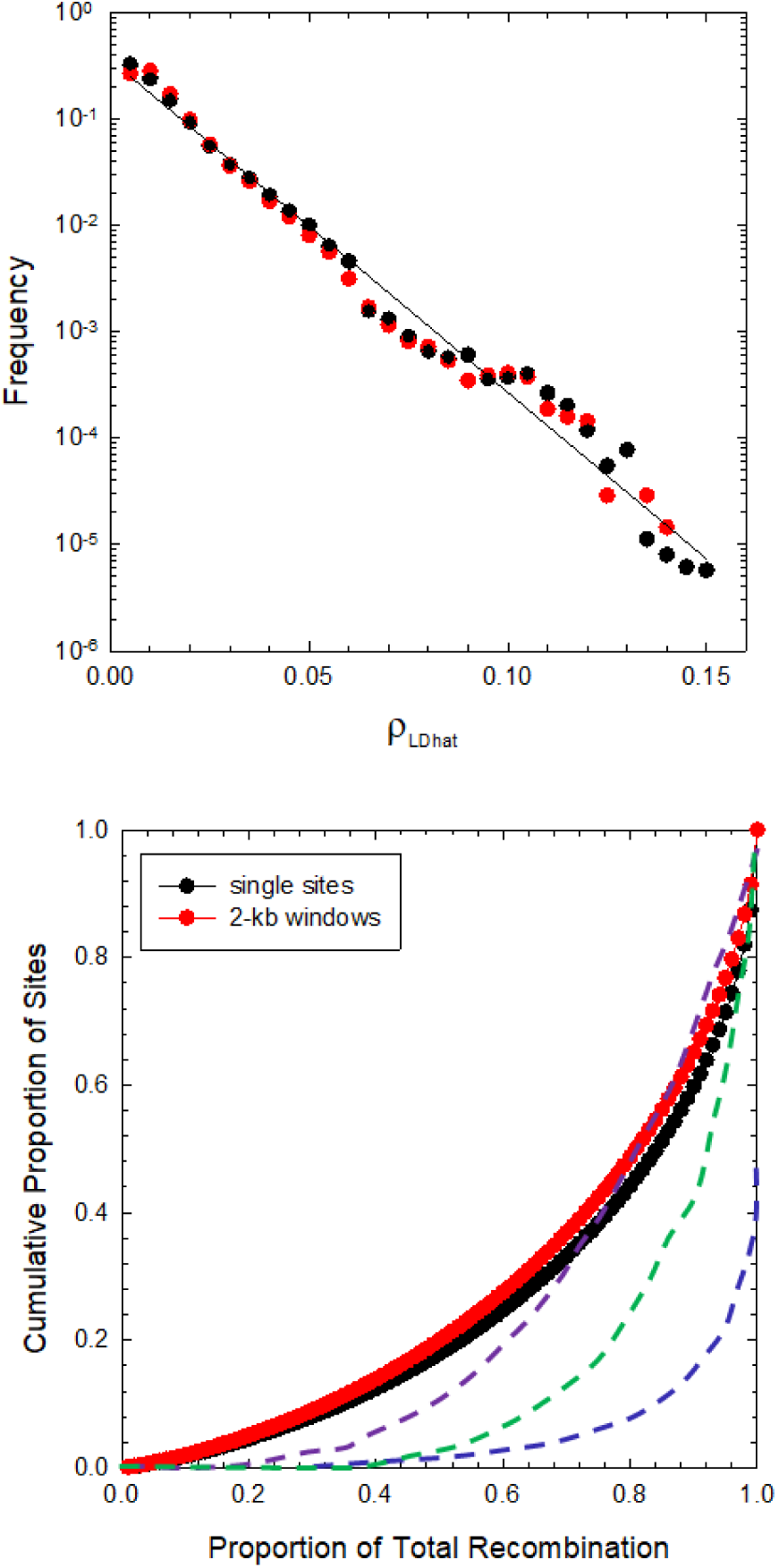
**Upper)** Frequency profiles of the 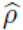 estimates of the population-averaged scaled recombination rate on a per-nucleotide and a per 2-kb window basis, based on 137,643,518 sites and 68,821 windows. The least-squares regression has intercept −0.60 and slope −33.0 (*r*^*2*^ = 0.979). **Lower**) Cumulative distributions for the data in the upper panel. The dashed blue line is the function derived for European human populations (taken from Figure 2 in Auton et al. 2012), green for mouse (Paigen et al. 2018), and purple for *Drosophila pseu doobscura* (Smukowski Heil et al. 2015).

To further evaluate whether a small subset of chromosomal regions do represent true recombination hotspots, we searched for such locations in each population sample using LDhot (Auton et al. 2014), using the output derived from LDhat as input files and simulating 1000 random data sets as null expectations. Although 536 hotspot regions (with average length 5995 bp and average inflation of the recombination rate of 21.5×) were identified by this procedure, the false-positive rate of inferring hotspots can be very high even in species containing legitimate hotspots (up to 50% false positives per Mb in human data; Auton et al. 2014), and it may be worse when the background recombination rate is only a few times lower than that in the potential hotspots (as in this study). An attempt was made to determine whether this subset of putatively high-recombination regions was enriched for specific nucleotide-sequence motifs by applying MEME 4.12.0 (Bailey et al. 2009). The only potential motif found was a largely homopolymeric run of Ts, but as this same motif is also enriched in non-hotspot regions, it remains doubtful that it is specifically an enhancer of the recombination rate.

To determine whether recombination rates are influenced by factors associated with gene bodies, we estimated how the average recombination rates per nucleotide site varied as a function of the distance from the known transcriptional-initiation sites in this species (Raborn et al. 2016). Although there is a general pattern of depressed recombination towards the beginning of genes (Figure 6), the magnitude of reduction is only ∼ 13% relative to the genome-wide average.

**Figure 6.**
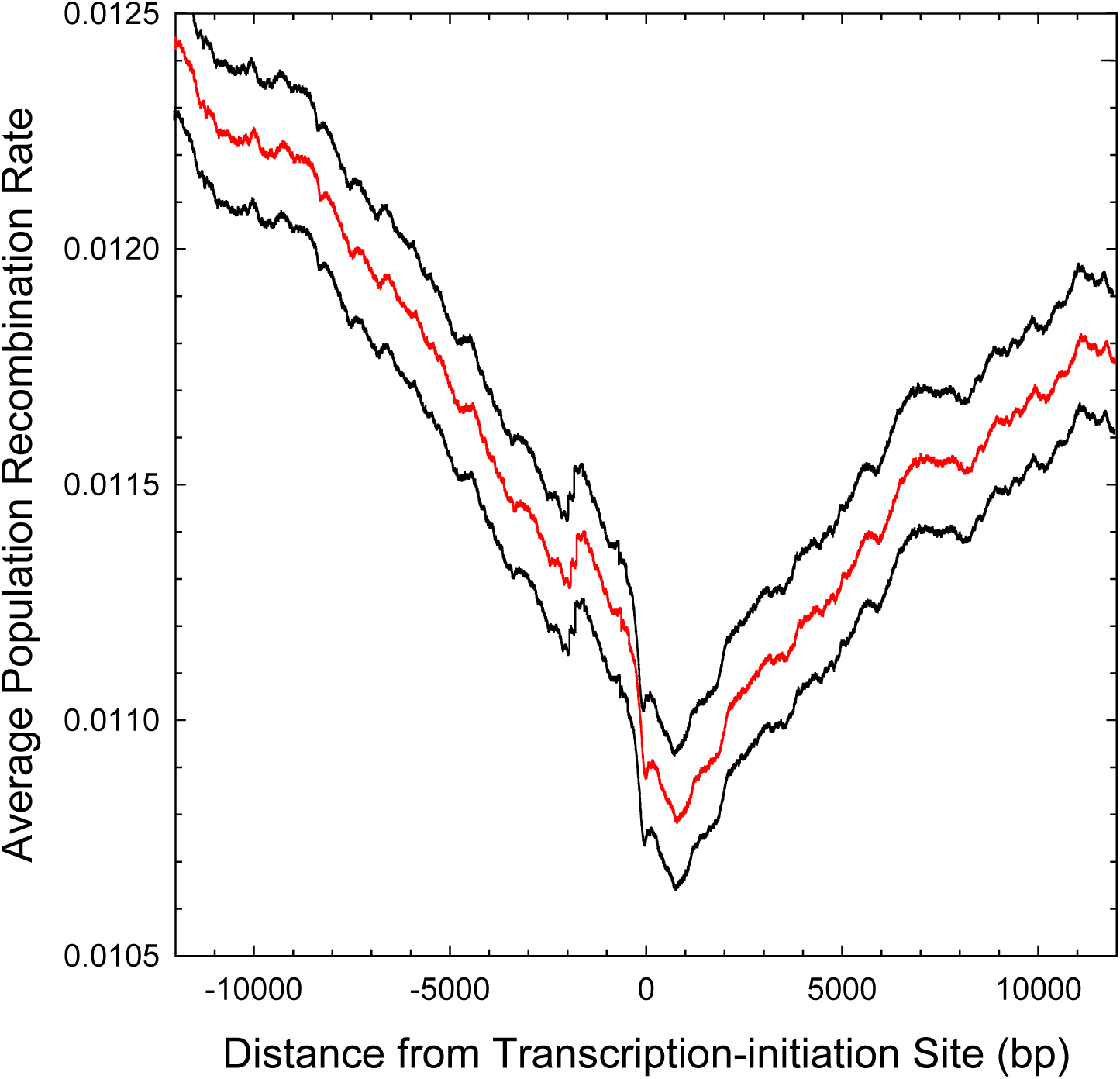
Average population-scaled recombination rate, as estimated by LDhat, as a function of the distance from the transcription-initiation sites of all genes for which such positions are known; the black lines denote ±2 standard errors of the means. The horizontal dashed line denotes the genome-wide average recombination rate.

As an alternative approach to revealing the recombinational landscape in a more sensitive way that accounts for the slow increase of the recombination rate with physical distance, we estimated *r*^2^ for all pairs of SNPs within sliding windows containing 101 polymorphic sites. To account for the mathematical constraints on *r*^2^, each estimate was scaled by dividing by the maximum possible value conditional on the allele frequencies at the two sites, thereby allowing all normalized estimates 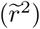 to fall in the full (0,1) range. Assuming a qualitative relationship of 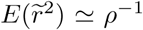, each pairwise 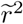 value was transformed into an estimate 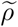 of 4*N*_*e*_*c* under the assumption of a −0.25 power-law scaling with distance (based on observations noted above), such that

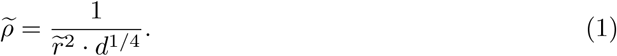

The full set of 100(100 − 1)*/*2 pairwise estimates within each window was then averaged to give window-wide estimates of 4*N*_*e*_*c*.

As can be seen in Figure 4, Equation (1) yields substantially higher estimates of *ρ* than does LDhat. While individual estimates are highly variable, when 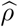 estimates are binned into windows, they are significantly correlated with average 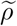 estimates obtained in the same intervals, ith the two scaling approximately linearly with respect to each other (Figure 7). Averaged over 65,837 windows, the mean site-specific estimate of 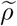 is 0.5376 (SE = 0.0005), 36× greater than mean 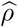, which is very similar to the expected 35-fold inflation noted above. Strikingly, the spatial variation in 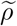 is greatly muted relative to that for 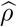 (Figure 4), further supporting the contention that *D. pulex* is lacking in recombination hotspots.

**Figure 7.**
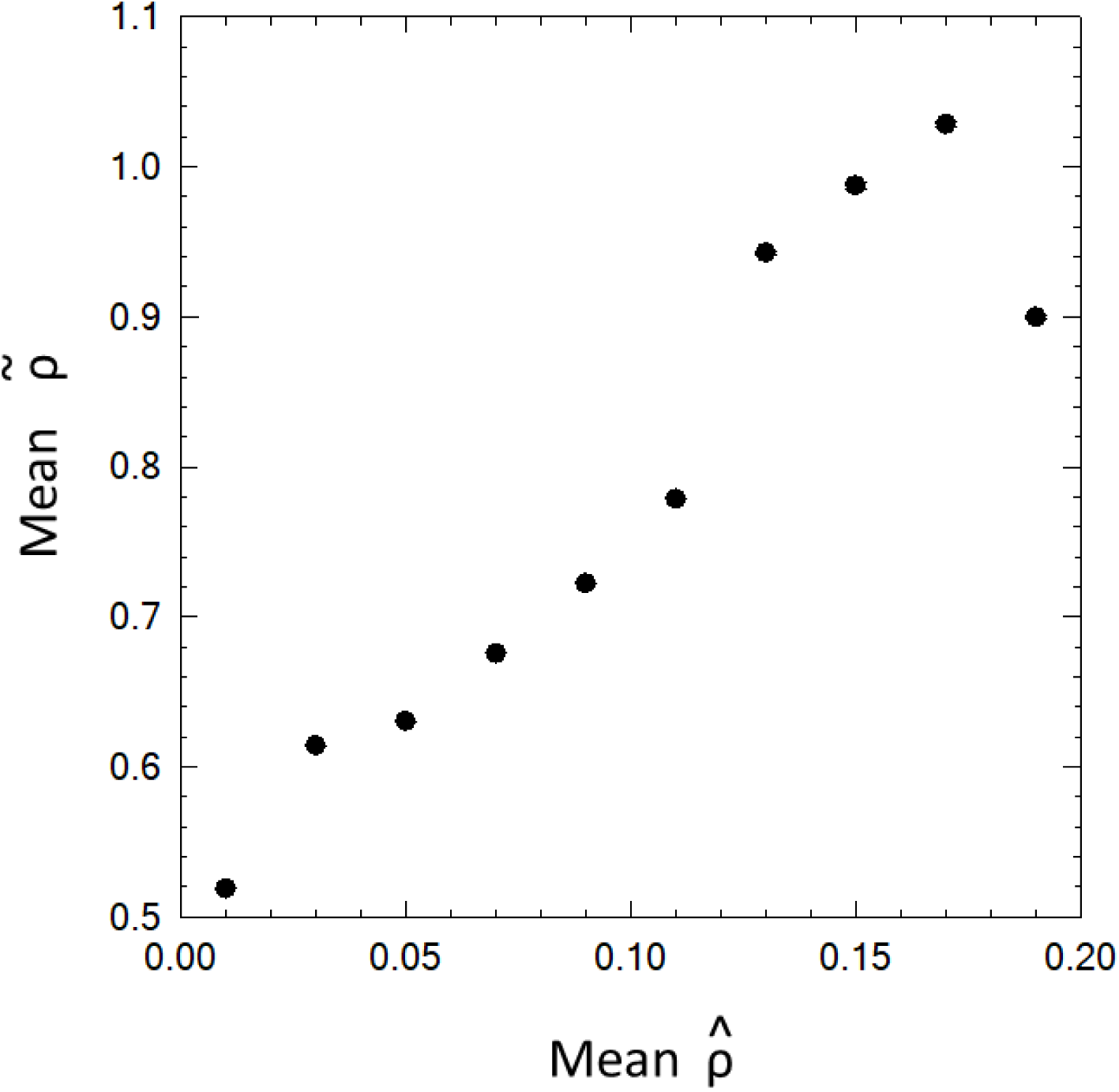
Average estimates of 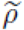 give n for 2-kb length regions binned on the basis of their 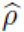 estimates. The standard errors of the former are smaller than the widths of the points on the graph.

By use of scaffolds that have been uniquely anchored to the genetic map (Xu et al. 2015), it was possible to estimate average recombination rates for each of the chromosomes (Table 2). For 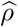, these fall in the narrow range of 0.0092 to 0.0157 (with average SE = 0.0013), and for 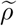 in the range of 0.499 to 0.616 (with average SE = 0.014). Owing to the typical eukaryotic pattern of ∼ 1 crossover per chromosome arm, one might expect the per-site recombination rate to decline with increasing chromosome length (Jensen-Seaman et al. 2004; Smeds et al. 2016; Stuckenbrock and Dutheil 2018). However, the estimated map lengths of chromosomes fall in the narrow range of 82 to 150 cM (Table 2), and chromosome-specific average 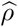 are uncorrelated with these estimates. The alternative statistic 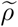 is negatively correlated with chromosome length at a significance level of 0.03 (*r*^2^ = 0.39), although the regression coefficient, −0.0096 (0.00039), is very weak.

**Table 2.** Estimates of average chromosome-specific 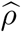 and 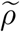, the two alternative measures of the per-site population recombination rate, based on the data from all scaffolds with known chromosomal locations. cM is the estimated length of the chromosome in centiMorgans (from Xu et al. 2015). Mb is the total length of chromosome-assigned scaffolds (in megabases), and this is followed by the number of scaffolds involved.

| Chromosome | cM | $\hat{\rho}$ | $\rho^*$ | Mb | Number |
| --- | --- | --- | --- | --- | --- |
| 1 | 141 | 0.0100 (0.0016) | 0.565 (0.015) | 8.6 | 17 |
| 2 | 150 | 0.0132 (0.0008) | 0.534 (0.011) | 11.6 | 33 |
| 3 | 116 | 0.0097 (0.0022) | 0.519 (0.020) | 8.5 | 21 |
| 4 | 82 | 0.0118 (0.0007) | 0.597 (0.011) | 4.0 | 9 |
| 5 | 133 | 0.0092 (0.0012) | 0.536 (0.016) | 11.6 | 31 |
| 6 | 119 | 0.0140 (0.0007) | 0.520 (0.014) | 6.5 | 19 |
| 7 | 139 | 0.0143 (0.0011) | 0.538 (0.013) | 9.9 | 30 |
| 8 | 113 | 0.0146 (0.0016) | 0.549 (0.014) | 10.4 | 26 |
| 9 | 119 | 0.0146 (0.0016) | 0.549 (0.019) | 8.4 | 26 |
| 10 | 139 | 0.0157 (0.0013) | 0.540 (0.012) | 13.3 | 37 |
| 11 | 83 | 0.0129 (0.0009) | 0.616 (0.007) | 4.8 | 10 |
| 12 | 121 | 0.0129 (0.0024) | 0.499 (0.018) | 8.5 | 15 |

Finally, there is a positive relationship between standing heterozygosity within nucleotide sites and the population-level recombination rate (Figure 8). Although the data are noisy for individual bins 2-kb in width, when the data are pooled into recombination-rate classes, it is clear that there is a positive relationship between these two parameters, with the first-order relationship being a scaling of average *π* with the 0.66 power of the measures of *ρ.*

**Figure 8.**
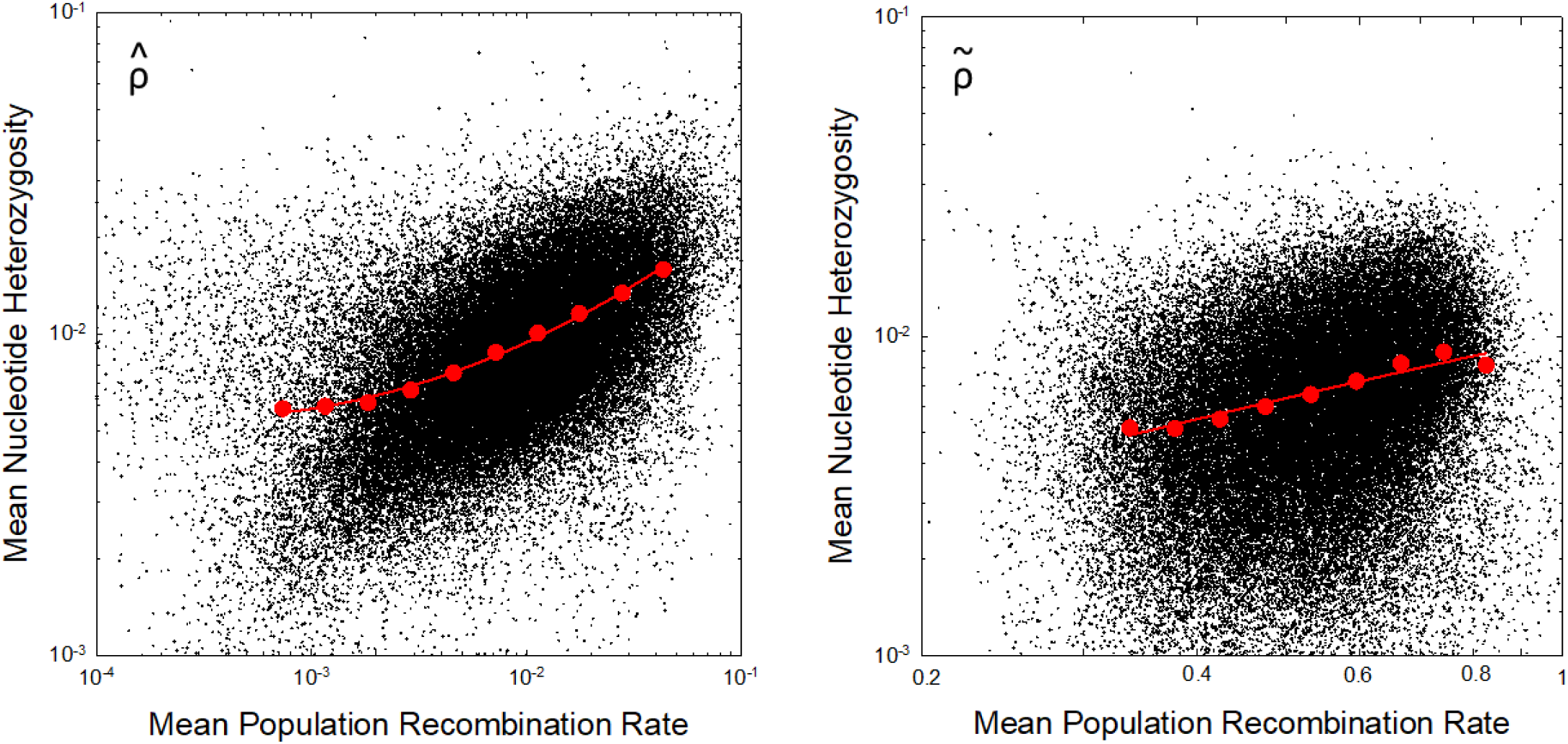
Relationship between average nucleotide heterozygosity and population recombination rate, 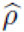, as estimated by LDHat. Small black points are based on windows of 2-kb width, whereas the larger red points are based on evenly binned data (on a logarithmic scale) for bins with sample sizes exceeding 1000 windows. The least-squares regression for the latter for 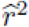 is 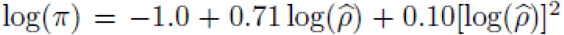, with *r*^*2*^ = 0.995, and for 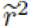 is 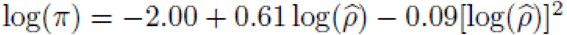, with *r*^*2*^ = 0.935.

## Discussion

This comprehensive study of *D. pulex* suggests a fairly homogeneous recombinational landscape in this species. Patterns of recombination as estimated from different populations are very similar, perhaps not surprisingly given the relatively low level of genetic subdivision among the sampled populations (Maruki et al., in prep.) Average recombination rates are similar among chromosomes, perhaps not surprisingly given the apparently similar physical lengths (Table 2). There is no compelling evidence of strong recombination hotspots, and although recombination appears to be suppressed in the vicinity of gene bodies, the effect is not large. Such suppression has been noted in *Drosophila* (Chan et al. 2012; Smukowski Heil et al. 2015) with a somewhat higher amplitude (20 to 50%). However, even here, we cannot entirely rule out intrinsic biases in the program used to estimate such rates, which might be associated with different levels of polymorphism (Dapper and Payseur 2018).

Because *Daphnia* generally undergo several generations of clonal propagation prior to engaging in sexual reproduction, it might be anticipated that they would have unusually high levels of linkage disequilibrium. However, relative to *D. melanogaster, D. pulex* actually experiences ∼ 60% more recombination per generation, even after accounting for periods of asexuality (Lynch et al. 2014). This result is largely a consequence of *Daphnia* having a similar genome sizes to that of *Drosophila*, but distributed over chromosomes (12 vs. 3), and of the latter lacking recombination in males. *D. pulex* and *D. melanogaster* have roughly comparable levels of LD for sites separated by fewer than 100 nucleotides (Lynch et al. 2014), but at larger distances LD is several-fold higher in *D. pulex* than in *D. melanogaster*, suggesting species differences in recombination mechanisms.

Although recombinational hotspots appear to account for very large fractions of the total recombinational activity in humans (Myers et al. 2005), chimpanzees (Auton et al. 2012), mouse (Paigen et al. 2008), and birds (Groenen et al. 2009; Kawakami et al. 2017), this appears not to be the case in *Daphnia.* Strong recombination hotspots are also lacking in *Drosophila* (Chan et al. 2012; Comeron et al. 2012; Manzano-Winkler et al. 2013; Smukowski Heil et al. 2015), although there does appear to be more heterogeneity in rates than in *D. pulex* (Figure 5). Kaur and Rockman (2014) also suggest an absence of hot spots in *C. elegans.* Thus, the current data suggest that, among animals, recombination hotspots may largely be confined to vertebrates.

Although many applications in population genetics assume that the rate of recombination increases linearly with the physical distance between sites, the results presented here, along with observations on a diversity of other species (Lynch et al. 2014, 2017), suggest that this assumption is far from fulfilled even at physical distances as large as 10^5^ kb. It remains unclear is why the scaling of LD with physical distance is so strong at short distances and so weak at long distances. Small-scale gene conversions along with the nonindependent arrival of mutational clusters are likely responsible for the rapid initial decline in LD with distance, contrary to an expected initial plateau followed by a simple downwardly bowed pattern if all recombination were simply due to crossing over in a linear distant-dependent pattern (Figure 3). However, the scaling at very large distances observed in Figures 1 and 2 are not easily reconciled without invoking a distribution of common and relatively long conversion-like events, which in principle could cause a prolonged intermediate shoulder in the LD-distance profile (Figure 3). Historical changes in population size can influence the distance-dependent pattern of LD (Lynch et al. 2014), but the data for *D. pulex* suggest historical changes of *N*_*e*_ by factors of no more than a few fold (Lynch et al. 2017, in prep.), which are inadequate to generate the patterns seen herein.

Given that the use of spatial patterns of LD to infer selection is a major enterprise (Walsh and Lynch 2018), the acquisition of unbiased inferences will require the development of proper null models based on an understanding of the recombinational mechanisms that generate baseline LD profiles. The quantitative relationships presented here provide a start, but these are purely statistical expressions, leaving the causal determinants unclear.

## Acknowledgments

This work was supported by NIH grants R01-GM101672 and R35-GM122566-01 and NSF grant DEB-1257806 to ML. We thank Ken Spitze and Emily Williams for help in sample collection and DNA preparation.

## Literature Cited

Andolfatto, P., and M. Nordborg. 1998 The effect of gene conversion on intralocus associations. Genetics 148: 1397–1399.

Auton, A., S. Myers, and G. McVean. 2014. Identifying recombination hotspots using population genetic data. arXiv preprint:1403.4264.

Auton, A., et al. 2012. A fine-scale chimpanzee genetic map from population sequencing. Science 336: 193–198.

Bailey, T. L., et al. 2009. MEME SUITE: tools for motif discovery and searching. Nucleic Acids Res. 37: W202–W208.

Chan, A. H., P. A. Jenkins, and Y. S. Song. 2012. Genome-wide fine-scale recombination rate variation in *Drosophila melanogaster.* PLoS Genet. 8: e1003090.

Charlesworth, B., and D. Charlesworth. 2010. Elements of Evolutionary Genetics. W. H. Freeman and Co., New York, NY.

Comeron, J. M., R. Ratnappan, and S. Bailin. 2012. The many landscapes of recombination in *Drosophila melanogaster.* PLoS Genet. 8: e1002905.

Dapper, A. L., and B. A. Payseur. 2018. Effects of demographic history on the detection of recombination hotspots from linkage disequilibrium. Mol. Biol. Evol. 35: 335–353.

Garud, N. R., and D. A. Petrov. 2016. Elevated linkage disequilibrium and signatures of soft wweeps are common in *Drosophila melanogaster.* Genetics 203: 863–880.

Groenen, M. A., et al. 2009. A high-density SNP-based linkage map of the chicken genome reveals sequence features correlated with recombination rate. Genome Res. 19: 510–519.

Haubold, B., P. Pfaffelhuber, and M. Lynch. 2010. mlRho – A program for estimating the population mutation and recombination rates from shotgun-sequenced genomes. Mol. Ecol. 19, Suppl. 1: 277–284.

Hill, W. G. 1975. Linkage disequilibrium among multiple neutral alleles produced by mutation in finite population. Theor. Pop. Biol. 8: 117–126.

Jensen-Seaman, M. I., T. S. Furey, B. A. Payseur, Y. Lu, K. M. Roskin, C. F. Chen, M. A. Thomas, D. Haussler, and H. J. Jacob. 2004. Comparative recombination rates in the rat, mouse, and human genomes. Genome Res. 14: 528–538.

Kaur, T., and M. V. Rockman. 2014. Crossover heterogeneity in the absence of hotspots in *Caenorhabditis elegans*. Genetics 196: 137–148.

Kawakami, T., C. F. Mugal, A. Suh, A. Nater, R. Burri, L. Smeds, and H. Ellegren. 2017. Whole-genome patterns of linkage disequilibrium across flycatcher populations clarify the causes and consequences of fine-scale recombination rate variation in birds. Mol. Ecol. 26: 4158–4172.

Keith, N., A. E. Tucker, C. E. Jackson, W. Sung, J. I. Lucas-Lledó, D. Schrider, S. Schaack, L. Dudycha, and M. Lynch. 2016. High mutational rates of large-scale duplication and deletion in *Daphnia pulex*. Genome Res. 26: 60–69.

Lynch, M. 2007. The Origins of Genome Architecture. Sinauer Assocs., Inc., Sunderland, MA.

Lynch, M. 2008. Estimation of nucleotide diversity, disequilibrium coefficients, and mutation rates from high-coverage genome-sequencing projects. Mol. Biol. Evol. 25: 2421–2431.

Lynch, M., M. Ackerman, K. Spitze, Z. Ye, and T. Maruki. 2017. Population genomics of *Daphnia pulex.* Genetics 206: 315–332.

Lynch, M., L.-M. Bobay, F. Catania, J.-F. Gout, and M. Rho. 2011. The repatterning of eukaryotic genomes by random genetic drift. Ann. Rev. Genomics Hum. Genet. 12: 347–366.

Lynch, M., S. Xu, T. Maruki, P. Pfaffelhuber, and B. Haubold. 2014. Genome-wide linkage-disequilibrium profiles from single individuals. Genetics 198: 269–281.

Manzano-Winkler, B., S. E. McGaugh, and M. A. Noor. 2013. How hot are drosophila hotspots? examining recombination rate variation and associations with nucleotide diversity, divergence, and maternal age in *Drosophila pseudoobscura.* PLoS One 8: e71582.

Maruki, T., and M. Lynch. 2014. Genome-wide estimation of linkage disequilibrium from population-level high-throughput sequencing data. Genetics 197: 1303–1313.

Maruki, T., and M. Lynch. 2015. Genotype-frequency estimation from high-throughput sequencing data. Genetics 201: 473–486.

Maruki, T., and M. Lynch. 2017. Genotype calling from population-genomic sequencing data. G3: Genes—Genomes—Genetics 7: 1393–1404.

McVean, G., and A. Auton. 2007. LDhat 2.1: A package for the population genetic analysis of recombination. Department of Statistics, Oxford, OX1 3TG, UK.

Myers, S., L. Bottolo, C. Freeman, G. McVean, and P. Donnelly. 2005. A fine-scale map of recombination rates and hotspots across the human genome. Science 310: 321–324.

Ohta, T., and M. Kimura. 1971. Linkage disequilibrium between two segregating nucleotide sites under the steady flux of mutations in a finite population. Genetics 68: 571–580.

Omilian, A. R., M. E. A. Cristescu, J. L. Dudycha, and M. Lynch. 2006. Ameiotic recombination in asexual lineages. Proc. Natl. Acad. Sci. USA 103: 18638–18643.

Paigen, K., J. P. Szatkiewicz, K. Sawyer, N. Leahy, E. D. Parvanov, S. H. Ng, J. H. Graber, W. Broman, and P. M. Petkov. 2008. The recombinational anatomy of a mouse chromosome. PLoS Genet. 4: e1000119.

Raborn, R. T., K. Spitze, V. P. Brendel, and M. Lynch. 2016. An atlas of promoters in the *Daphnia* genome revealed by comprehensive mapping of 5I-mRNA ends. Genetics 204: 593–612.

Smeds, L., C. F. Mugal, A. Qvarnström, and H. Ellegren. 2016. High-resolution mapping of crossover and non-crossover recombination events by whole-genome re-sequencing of an avian pedigree. PLoS Genet. 12: e1006044.

Smukowski Heil, C. S., C. Ellison, M. Dubin, and M. A. Noor. 2015. Recombining without hotspots: a comprehensive evolutionary portrait of recombination in two closely related species of *Drosophila.* Genome Biol. Evol. 7: 2829–2842.

Stukenbrock, E. H., and J. Y. Dutheil. 2018. Fine-scale recombination maps of fungal plant pathogens reveal dynamic recombination landscapes and intragenic hotspots. Genetics 208: 1209–1229.

VanLiere, J. M., and N. A. Rosenberg. 2008. Mathematical properties of the *r*^2^ measure of linkage disequilibrium. Theor. Popul. Biol. 74: 130–137.

Walsh, J. B., and M. Lynch. 2018. Evolution and Selection of Quantitative Traits. Oxford Univ. Press, UK.

Xu, S., M. S. Ackerman, H. Long, L. Bright, K. Spitze, J. S. Ramsdell, W. K. Thomas, and M. Lynch. 2015. A male-specific genetic map of the microcrustacean *Daphnia pulex* based on single sperm whole-genome sequencing. Genetics 201: 31–38.

Ye, Z., S. Xu, K. Spitze, J. Asselman, X. Jiang, M. S. Ackerman, J. Lopez, B. Harker, R. T. Raborn, M. E. Pfrender, and M. Lynch. 2017. Comparative genomics of the *Daphnia pulex* species complex. G3: Genes, Genomes, Genetics 7: 1405–1416.

